# A Regulatory Role of Molecular Chaperones in Excited States and Functions of Folded Enzymes

**DOI:** 10.1101/2025.07.15.664663

**Authors:** Guan Wang, Bin Yu, Yihao Chen, Ruishen Ding, Shixing Qi, Xiaoling Zhao, Zhou Gong, Xu Zhang, Sebastian Hiller, Maili Liu, Lichun He

## Abstract

Molecular chaperones are recognized for assisting protein folding. Emerging evidence suggests that chaperones also interact with natively folded proteins. Yet their functional impact on folded proteins remains unclear. Here, we show that chaperones can directly modulate the conformational dynamics and catalytic efficiency of enzymes in their native states. Spy and Hsp70 increased lysozyme activity by altering conformational exchanges, and this effect extended across other chaperones and enzymes. Hsp70, Spy, Hsp104, and Hsp20 enhanced the catalytic activities of multiple folded enzymes, including alkaline phosphatase, DNA polymerase Pfu, endonucleases Cas12a/Cas13a, and xylanase. These findings uncover an unanticipated function of chaperones in regulating enzyme function expanding their mechanistic scope beyond folding assistance and suggests opportunities in enzyme engineering, diagnostics, and cellular regulation.

## Main Text

Molecular chaperones, comprising about 10% of cellular proteins, play a critical role in protein quality control (*1, 2*), such as protein folding, translocation, unfolding, disaggregation, and maintaining the homeostasis of client proteins. In past decades, many studies have revealed mechanisms about chaperones in aiding protein folding, that chaperones interact with nascent polypeptides during synthesis, shielding hydrophobic regions to prevent misfolding and aggregation (*3*). Additionally, chaperones interact with misfolded proteins to induce destabilization and facilitate their proper refolding (*4*). However, growing evidence suggests that molecular chaperones also interact with natively folded proteins. For example, Hsp70 and Hsp90 chaperones associate with 60% of kinases, 30% of ubiquitin ligases, and 7% of transcription factors (*5*). Chaperones are found not only in the cytoplasm but also in the nucleus, where no nascent polypeptide translation occurs (*6-8*). Hsp70s were reported to accumulate in the nucleus in response to heat stress where no nascent polypeptide synthesis occurs (*9*). 2-3% of the cellular Hsp90 were pooled in the nucleus (*10, 11*). Thus, it is critical to know whether and how chaperones played roles beyond its traditional functions of aiding protein folding and preventing protein aggregation.

In our previous work, we investigated the recognition and binding mechanisms of chaperones with folded client protein. Different chaperones Spy, SurA and Skp recognized and interacted with the common frustrated surface of folded client proteins Im7 and Fyn SH3 (*12, 13*). Moreover, serum albumin which was also reported to display chaperone-like activity in the extracellular environment by associated with client proteins (*14-16*). Bovine serum albumin (BSA) was revealed to alter the folding energy landscape of Fyn SH3 domain and induced a new excited state (*17*). The excited state of a proteins is a transient, higher-energy conformation that protein adopts due to thermal fluctuations or external perturbations. The excited state often has functional significance, enabling proteins to carry out enzymatic catalysis, allosteric regulation, and molecular recognition. Regarding to enzymes, they usually adopted multiple conformations with the equilibrium of a ground state and one or more excited states. Each excited state may display a distinct level of activity (*18*). Many studies showed the changes of conformations of enzymes altered the activities of enzymes (*19, 20*). As an enzyme transitions between conformations, its substrate affinity and catalytic efficiency fluctuate, reflecting shifts in its ability to convert substrate into product. The range of conformational states available to an enzyme is dictated by its energy landscape. Various factors can reshape energy landscape, subsequently affecting enzymatic activity (*21*). Given that chaperones have been shown to modulate the folding energy landscape by stabilizing or destabilizing specific conformations of client proteins (*22, 23*), we speculate that they may also have the potential to influence the function of folded enzymes.

To investigate this new role of chaperones, we evaluated the impact of molecular chaperones on the catalytic activity of six naturally folded enzymes, including hen egg-white lysozyme (HEWL), alkaline phosphatase PHPT1, DNA polymerase from *Pyrococcus furiosus* (*Pfu* pol), endonuclease Cas12a and Cas13a, xylanase. Four chaperones from different categories such as *Homo sapiens* Hsp70 (A8), *Thermus thermophilus* Hsp104 (ClpB), *Escherichia coli* chaperone Spy (Spy), and *Thermococcus Kodakaraensis* Hsp20 (TkHsp20) were chosen as additives to investigate its impact on reactions catalyzed by distinct enzymes respectively. The results showed molecular chaperones enhanced the catalytic activities of all six investigated enzymes. Further experiments via NMR spectroscopy revealed that the molecular chaperones Spy and Hsp70 A8 enhanced HEWL activity by modulating its conformational equilibrium without altering its structure. This study reveals an unexpected role of chaperones in modulating the activities of folded enzymes, underscores their potential for enhancing enzyme performance in diagnostics and biomanufacturing, and sheds light on emerging functions of upregulated chaperones in orchestrating systemic modulation of cellular processes.

## Results

### Molecular chaperones enhance the activity of folded HEWL

To investigate whether molecular chaperones can interact with folded enzymes and modulate their activity, we firstly selected HEWL as a model client. HEWL consists of 129 amino acid residues and features a structure of six α-helices and a three-stranded β-sheet. It exerts bacteriolytic activity by hydrolyzing β-1,4-glycosidic bonds within bacterial peptidoglycan. Addition of 0.7 μM HEWL to *Micrococcus lysodeikticus* (1.2 mg/mL) led to a marked decrease in absorbance of light with a wavelength of 450 nm (OD450), confirming enzymatic lysis of *Micrococcus lysodeikticus* (Fig. 1A), whereas the control group without HEWL showed no change of OD450 absorbance. We next evaluated the impact of four different molecular chaperones Hsp70 A8, ClpB, Spy, and TkHsp20 on the activity of HEWL. All four chaperones enhanced HEWL activity, with distinct potency (Fig. 1A), while chaperones alone showed no hydrolytic activity, as no significant changes of OD450 were observed over time (Fig. S1B). Notably, the activity enhancement of HEWL by chaperones was in a dose dependent manner (Fig. 1D, Fig. S2). HEWL activity was progressively enhanced with increasing concentrations of molecular chaperones. Spy increased HEWL activity from 1.3-fold to 2.4-fold when the molar ratio of Spy and HEWL increased from 1:1 to 6:1 (Fig. 1B). Similarly, Hsp70 A8 achieved a boost of HEWL activity from 1.2-fold to 2.2-fold when the molar ratio of Hsp70 A8 and HEWL increased from 0.05:1 to 0.5:1 (Fig. 1C). These results demonstrated that molecular chaperones, though catalytically inert, could potentiate the enzymatic activity of a folded client protein, with variable efficacy depending on chaperone types and concentrations.

**Figure 1.**
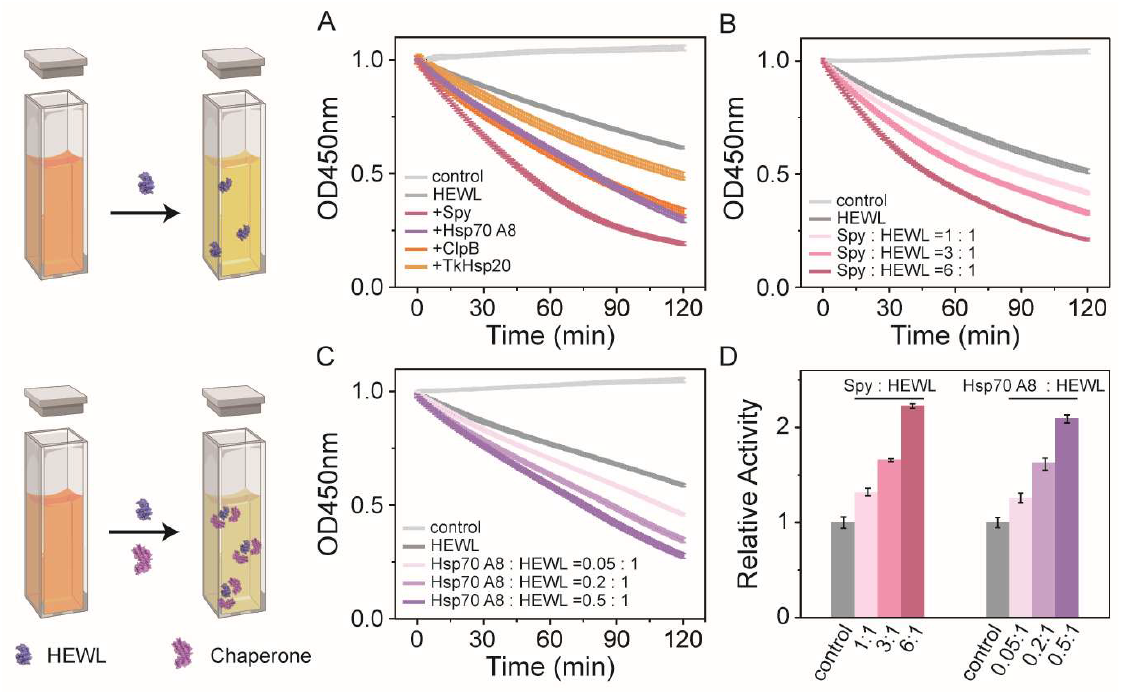
Activity assays of HEWL in the presence of molecular chaperones. (A) Time-dependent enzymatic activity assays of HEWL (PDB 2EPE) in the presence of molecular chaperones Spy, Hsp70 A8, ClpB, and TkHsp20 at molar radios of 6:1, 0.5:1, 5:1, and 5:1 relative to HEWL, respectively. (B) Time-dependent enzymatic activity assays of HEWL in the presence of Spy at molar radios of 1:1, 3:1, and 6:1 relative to HEWL. (C) Time-dependent enzymatic activity assays of HEWL in the presence of Hsp70 A8 at molar radios of 0.05:1, 0.2:1, and 0.5:1 relative to HEWL. The grey line in both panels corresponds to the control experiment, which is performed in the absence of HEWL in the cell suspension. (D) The relative activity assays of HEWL in the presence of different concentrations of chaperones Spy and Hsp70 A8, normalized to HEWL alone (control). Data represent the mean ± SD of three technical replicates.

## Molecular chaperones Spy and Hsp70 A8 interact with HEWL in a non-specific and transient way

In a next step, we employed solution NMR spectroscopy and isothermal titration calorimetry (ITC) experiment to examine the interaction between HEWL and chaperones to understand how molecular chaperones enhance the activity of HEWL. The ITC experiment revealed HEWL interacted with chaperone Spy with dissociation constant (*K*_D_) in the range of millimolar (Fig. S3). ^15^N-labeled HEWL was expressed in *P. pastoris* according to the protocol mentioned in the method and material section. 2D [^15^N,^1^H]-heteronuclear single quantum coherence (HSQC) spectra of 1 mM ^15^N-labeled HEWL were recorded in the presence and absence of different concentrations of Spy and Hsp70 A8 respectively (Fig. 2A, Fig. S5A). Strikingly, the spectra overlapped closely. The chemical shift perturbation of all residues was less than 0.05 ppm, indicating the conformation of HEWL was not perturbed upon adding either the chaperone Spy or the chaperone Hsp70 A8 (Fig. S4A and Fig. S5B). However, peak intensities of ^15^N HEWL dropped—by 43% with equimolar Spy and by 53% with Hsp70 A8 at 0.05:1 molar ratio (Fig. 2B, Fig. S5C). The plot of intensity ratio of the peaks from HEWL in the presence and absence of the chaperone Spy or Hsp70 A8 against its sequence revealed almost every residue of HEWL showed similar peaks intensity reduction, suggesting HEWL interacted with Spy or Hsp70 A8 in a global and non-specific manner. Furthermore, we performed the solvent paramagnetic relaxation enhancement (PRE) experiment. 2 mM Gd-DOTA was added in the NMR buffer to serve as the solvent PRE reagent, which could bleach the NMR signal of HEWL residues exposed to the solvent. If stable complex between Spy and HEWL formed, NMR signals of HEWL residues from the interface should be protected from bleaching by the chaperone Spy (*13*). Thus, we determined the shielding index (σ_i_) of Spy on HEWL according to a previously published method (*13*). Intriguingly, no cluster of residues of HEWL protected by Spy was observed (Fig. 2C), indicating HEWL interacted with Spy and Hsp70 A8 in a transient and fuzzy way.

**Figure 2.**
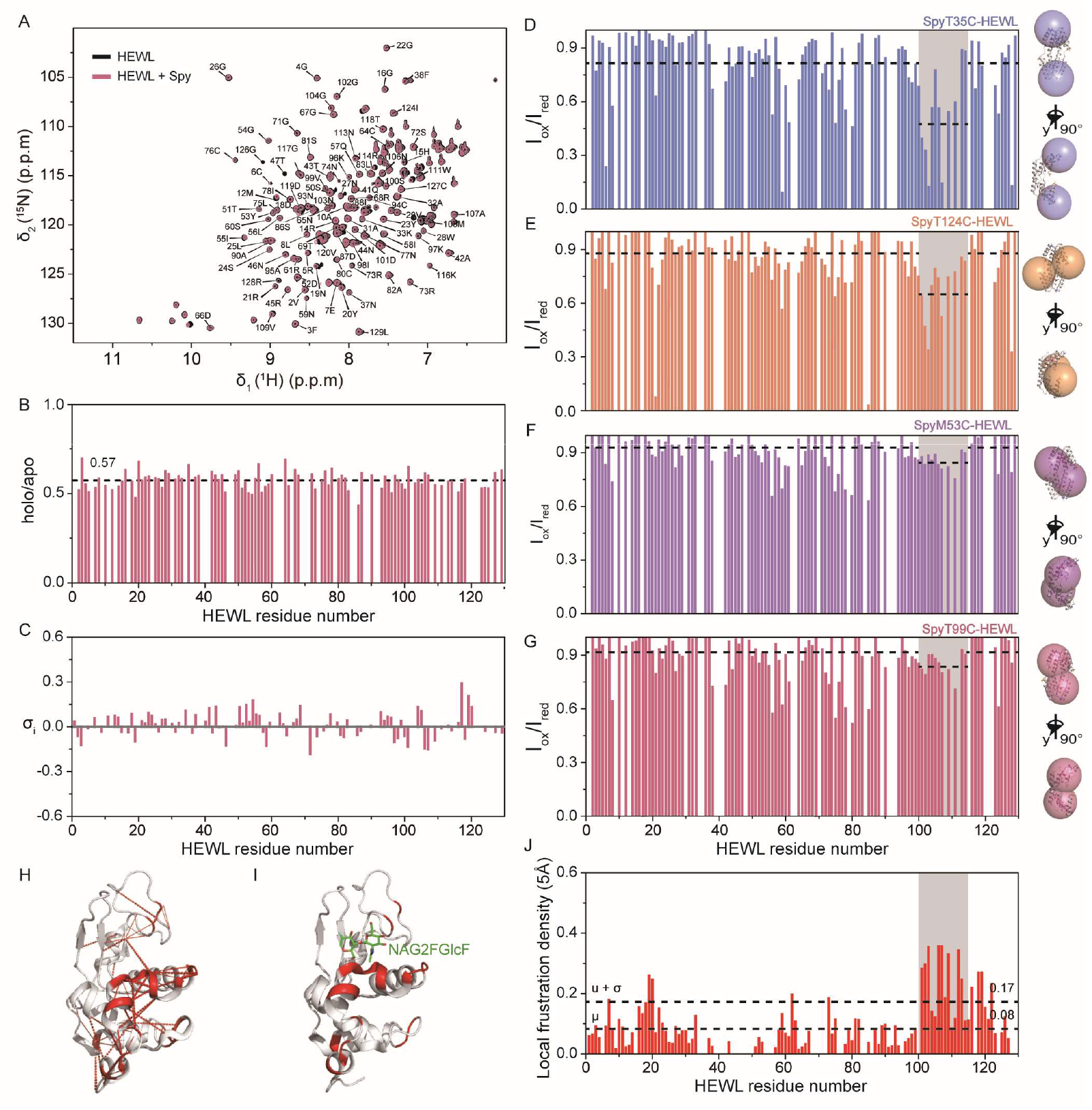
Molecular chaperone Spy interact with HEWL in a non-specific and transient way. (A) Overlay of [^15^N,^1^H]-HSQC spectra of 1 mM HEWL in the absence (black) and presence (red) of 1 mM Spy. (B) Ratios of the residual resolved cross peak volume of backbone amides of 1 mM ^15^N-labeled HEWL in the presence and absence of 1 mM Spy plotted against the corresponding amino acid residue number. (C) Solvent PRE shielding effect index σ_i_ of Spy on ^15^N-labeled HEWL plotted against the corresponding amino acid residue number. (D-G) Intermolecular paramagnetic relaxation effect on ^15^N-labeled HEWL upon interaction with the spin label MTSL-attached T35C (D), T124C (E), M53C(F) and T99C (G) mutant of Spy respectively. I_ox_/I_red_ indicates the ratio of peak heights of HEWL before and after the reduction of the MTSL spin label. (H-J) The local frustration of HEWL was calculated via the online web server http://frustratometer.qb.fcen.uba.ar (*25*). Distribution of highly local frustration was projected in amino acid sequence (J). Residues with local frustration density values larger than the averaged values plus standard deviation were depicted on the crystal structure of HEWL (PDB 2EPE) in red. Clusters of the highly frustrated contacts are colored red with red dashed lines (H). Superimposing of the highly frustrated residues onto the crystal structure of the E35Q-substrate complex (PDB 1H6M). The substrate NAG2FGlcF was depicted in green (I).

To further investigate this non-specific and transient interaction way, intermolecular PRE experiments were performed. As Spy does not contain any free cysteine, residue T35, M53, T99, and T124 of Spy were mutated to cysteine and labeled with (1-oxyl-2,2,5,5-tetramethylpyrroline-3-methyl) methanethiosulfonate (MTSL) individually. These four mutations introduced are broadly and evenly scattered over the surface of Spy, encompassing the entire protein (Fig. S4C, D). The labeled MTSL can reduce NMR signals of atoms within a radius of ∼25 Å of the mutated residue (*24*). Typically, this intermolecular PRE effect is strong within 20 Å and significant signal broadening occurs due to dipolar interactions with the unpaired electron of MTSL (*12, 24*). We plotted spheres with a radius of 20 Å centered in each of the four mutated residue T35, M53, T99, and T124 on the crystal structure of Spy (PDB 3O39). The spheres covered the whole surface of Spy (Fig. S4C, D). Given HEWL formed a stable and strong complex with Spy via any interface, it would fall into the strong bleaching radius of 20 Å. Broadening and intensity reduction of peaks from residues on the contact surface of HEWL would be observed. Yet again, no significant peak intensity decline was observed for HEWL upon the addition of any of the four MTSL labeled Spy variants individually (Fig. 2D-G), confirming that the interaction between Spy and HEWL was in a transient or weak way.

This transient and weak interaction way was also consistent with the negligible significant chemical shift perturbation of HEWL upon adding chaperones, since the chemical shift perturbation was either too weak or averaged out due to the different transient binding ways between HEWL and chaperones. Nevertheless, the region from residue 100 to 112 of HEWL showed a common and slight bleaching pattern upon adding all four differently MTSL labeled Spy, implying this region plays an important role for the weak and transient interaction between Spy and HEWL (Fig. 2D-G). Remarkably, this region from residue 100 to 112 was also the maximum local frustration region of HEWL (Fig. 2H, J). Frustration sites are localized regions within a protein where competing or conflicting interactions between residues hinder the attainment of a global energy minimum. These regions are frequently located at enzyme active sites and protein– protein interaction interfaces. This result was consistent with our previous study that chaperone recognizes the locally frustrated surface of the client proteins and that the interaction is transient and highly dynamic with no stable complex formed, leaving the chaperone-interacting surface solvent accessible (*13*). This transient interaction explained although the region 100 to 112 is close to the substrate binding site (Fig. 2I), the access of the substrate to HEWL was not blocked and the catalytic efficiency of HEWL in the presence of chaperone Spy and Hsp70 A8 was increased more than 2 times.

### Molecular chaperones enhance the activity of HEWL through modulating its conformational exchanges

To gain insight into the mechanism chaperones enhance the activity of HEWL. We firstly measured if the chaperone changed the dynamics of HEWL, as the function of an enzyme could be influenced by its dynamics (*26-28*). Faster motions, occurring within picoseconds to nanoseconds, can generate significant conformational entropy, influencing binding and allosteric interactions (*29-31*). Therefore, T_1_, T_2_ and ^15^N-{^1^H} heteronuclear NOE of HEWL in the presence and absence were measured. The residue-specific spectral density function J(ω), which characterizes molecular dynamics across various time scales, is also determined at discrete frequency points based on measurements of NMR spin relaxation parameters (*32*) (Fig. S6). ^15^N-{^1^H} heteronuclear NOE and J (0.87ω_H_) showed the fast local protein backbone dynamics in the ps-ns timescale was not changed significantly (Fig. S6). Further the H-D exchange rate of HEWL measured by NMR spectroscopy revealed that the backbone amide proton exchanging free energy of HEWL in the absence and presence of 2 mM Spy had an average value of 6.52 and 6.69 kcal/mol respectively (Fig. S4B), indicating a highly similar folding status of HEWL in the presence and absence of Spy. The structural protection and intrinsic exchange rates of individual residues of HEWL was not perturbed by adding 2 mM Spy.

Furthermore, ^15^N Carr-Purcell-Meiboom-Gill relaxation dispersion (CPMG-RD) experiments were performed to assess the proportion of ground-state and excited-state conformations of HEWL in the presence or absence of molecular chaperones Spy and Hsp70 A8 at two different magnetic fields, 600 MHz and 800 MHz respectively (Fig. 3B). CPMG-RD is sensitive to molecular motions in proteins occurring on the microsecond-to-millisecond (μs-ms) timescale, which typically corresponds to conformational changes and allosteric transitions. 33 residues of HEWL exhibited non-flat dispersion curves indicating significant conformational exchanges were taking place (Fig. S7, Fig. S8). To compare the differences of conformational exchanges of HEWL in the apo- and holo-form with chaperone Spy and Hsp70 A8, we used the parameter ΔR_2,eff_ to indicate the chemical exchange differences, where ΔR_2,eff_ was calculated as the subtraction of {R_2,eff_ (ν_CPMG_ = 50 Hz) by R_2,eff_ (ν_CPMG_ = 1000 Hz)}. The large absolute value of ΔR_2,eff_ of HEWL in its apo- or holo-form implied the persisting dynamic conformational exchanges of HEWL. Notably, a similar region consisting of β strand 1, β strand 3, helix 4, along with the loop in between of β strand 1 and helix 4, and N-terminal loop on HEWL (Fig. 3H, I) undergoing conformational exchanges was observed for HEWL in both apo- and holo-form. This result was confirmed by CEST NMR experiments, which could directly demonstrate the chemical shifts and numbers of different conformations on a residue resolution level.

**Fig 3.**
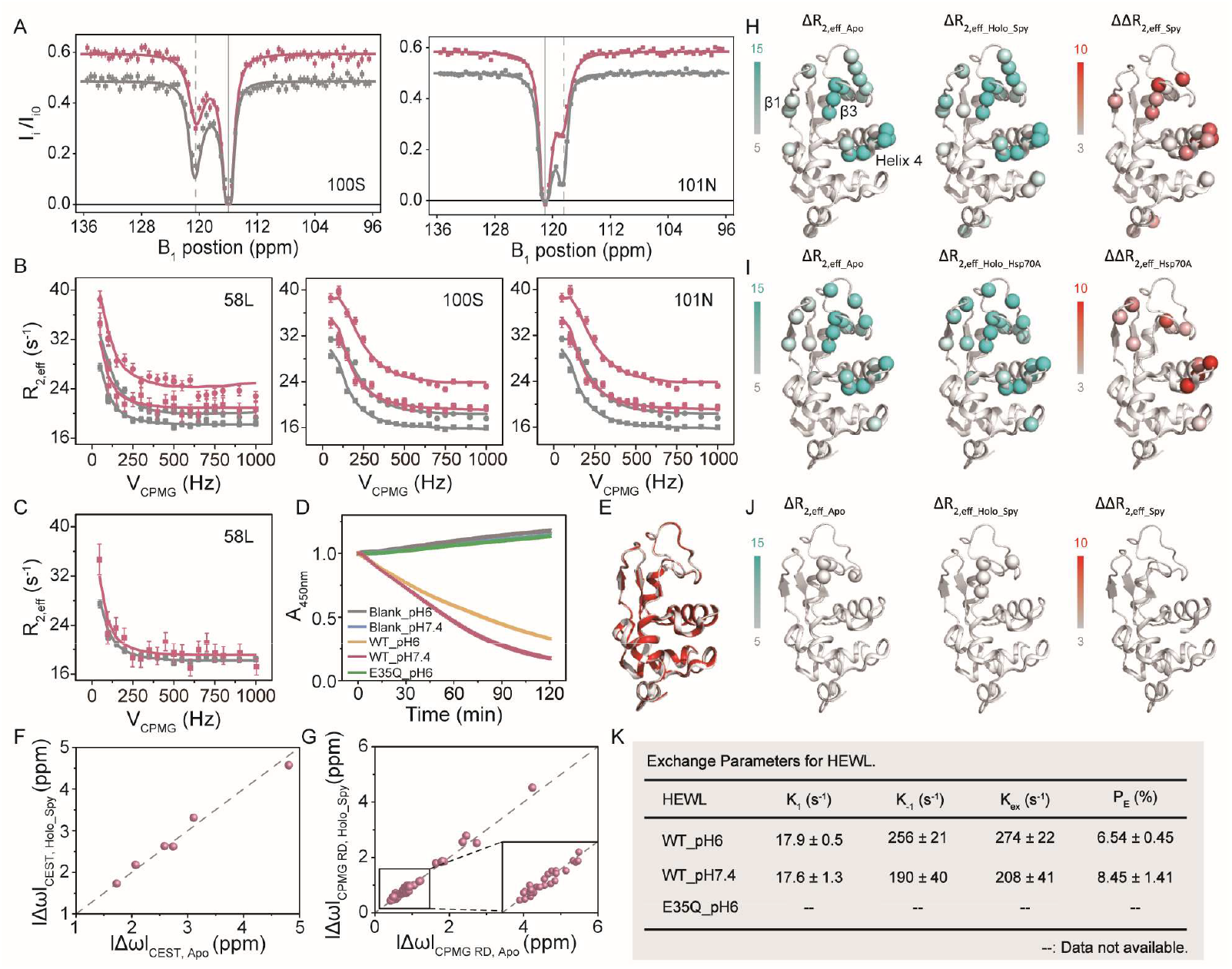
Molecular chaperones increase the proportion of the excited state conformation of HEWL. (A) Representative ^15^N CEST profiles of HEWL in the absence (gray) and in the presence (red) of the molecular chaperone Spy, obtained as described in the Methods and Materials at ^1^H frequencies of 600 MHz. (B) Representative ^15^N CPMG RD profiles of HEWL in the presence (red) and absence (gray) of the molecular chaperone Spy, obtained as described in the Methods and Materials at ^1^H frequencies of 600 (circle) and 800MHz (square). (C) Representative ^15^N CPMG RD profiles of HEWL in the presence (red) and absence (gray) of the molecular chaperone Hsp70 A8, obtained as described in the Methods and Materials at ^1^H frequencies of 600 MHz. (D) Time-dependent enzymatic activity of wide type and E35Q mutant of HEWL in different buffers with pH 6.0 and pH 7.4 respectively. (E) Structural align of HEWL (PDB 2EPE) with the E35Q mutant (PDB 3OK0), RMSD = 0.15 Å. (F) Correlation between the absolute |Δω| values for HEWL in the presence and absence of the molecular chaperone Spy extracted from CEST experiments. (G) Correlation between the absolute |Δω| values for HEWL in the presence and absence of the molecular chaperone Spy extracted from CPMG RD experiments. (H-J) Residues with significant ΔR_2_ and ΔΔR_2_ of backbone amides of 1 mM HEWL in the absence and presence of 2 mM Spy (H) and 0.05mM Hsp70 A8 (I) are shown as spheres and mapped onto the crystal structure of HEWL. ΔR_2_ and differences in ΔR_2_ (ΔΔR_2_) of backbone amides of E35Q mutant in the absence and presence of 2 mM Spy are mapped onto the crystal structure of E35Q mutant (J). A gray to blue color scale represents the ΔR_2_ value from 5 (gray) to 15 (blue), while a gray to red color scale represents the ΔΔR_2_ value from 3 (gray) to 10 (red). (K) Conformational exchange parameters for HEWL fitted from the CPMG RD data obtained in different buffers with pH 6.0 and pH 7.4 respectively.

^15^N CEST data indicated that HEWL undergoes two-state conformational exchanges in both apo- and holo-form (Fig. 3A and Fig. S9, S10). Moreover, ΔΔR_2,eff_ was introduced as the subtraction of ΔR_2,eff_ of apo-HEWL from ΔR_2,eff_ of holo-HEWL with chaperone Spy or Hsp70 A8 to indicate the difference of the dynamic conformational exchanges. Residues from parts of β strand 1, β strand 3 and helix 4, along with the loop in between of β strand 2 and helix 4 of HEWL, which were around the substrate binding site, undergo significantly different conformational exchanges in its holo-from with either Spy or Hsp70 A8 comparing to that in its apo-form (Fig. 3H, I). Following this up, a global fit using a two-state model was applied to these residues exhibiting significant chemical exchanges to determine the chemical shifts, populations, and exchange rates of HEWL in different states. Both chaperones, Spy and Hsp70 A8, influenced the thermodynamic equilibrium and kinetic properties of HEWL’s conformational changes. Specifically, the presence of 5% Hsp70 A8 (relative to HEWL) increased the population of HEWL’s excited state (P_E_) from 8.44 ± 1.43% to 14.52 ± 1.62%, while the presence of 50% Spy (relative to HEWL) caused a shift in P_E_ of HEWL from 6.53 ± 0.45% to 7.42 ± 0.37%. (Table 1). The conformational exchanges of apo-state of HEWL were measured in different buffer with pH 6.0 and pH 7.4 for comparing with its corresponding holo-state with Spy (pH 6.0) and Hsp70 A8 (pH 7.4) respectively. As the complex of Hsp70 A8 with HEWL were not stable in buffer with pH 6. Nevertheless, HEWL in pH 7.4, presented a larger population of the excited state and showed a higher activity than HEWL in pH 6.0, indicating the excited state of HEWL was related to its activity (Fig. 3C, D, K). Moreover, as the concentration of the molecular chaperone Spy increased, the population of excited-state conformations of HEWL increased. At a molar ratio of 2:1 (relative to HEWL), Spy shifted the population of the excited state of HEWL to 8.81 ± 0.86%. (Fig. 3B and Table S1). This result was in line with the observation that molecular chaperones enhance the bacteriolytic activity of HEWL in a concentration-dependent manner (Fig. 1D). In contrast, CPMG-RD experiments of the E35Q mutant of HEWL which abolished the bacteriolytic activity revealed it exhibited no excited-state (Fig. 3J). The addition of equal molar of Spy induced neither bacteriolytic activity nor emerging of the excited state of E35Q mutant (Fig. S1A, Table S1). The ΔR_2,eff_ values of HEWL E35Q mutant were close to zero in both the presence and absence of Spy. Remarkably, the structure of E35Q mutant was almost the same as the ground state structure of wide type HEWL, with a RMSD value of 0.15 (Fig. 3E), substantiating that the conformation exchanged to the excited-state of HEWL is essential for its activity. Additionally, the value of chemical shifts of the excited state measured for wide type HEWL in its apo- and holo-form with chaperones by both CEST and CMPG RD experiments correlated very well, indicating the excited-state structure of HEWL remained unchanged upon interacted with chaperones (Fig. 3F-G). Chaperones enhanced the bacteriolytic activity of HEWL by modulating its dynamics including the conformational exchanging rate and populations of different states.

### Molecular chaperones modulate activities of various enzymes

Since most enzymes are highly dynamic, their functions rely on conformational changes to convert substrates. Chaperones inherently possess the ability to monitor and modulate protein structures. This led us to investigate whether the activity-enhancing effect of chaperones on HEWL also applies to other enzymes. To explore this, we examined the activities of five additional enzymes with diverse enzymatic roles and applications in biology, including PHPT1, *Pfu* pol, Cas12a, Cas13a, and xylanase, in the presence of various chaperones from different categories. PHPT1 is an enzyme involved in the signal transduction in cells. It hydrolyzes its substrate, para-nitrophenyl phosphate (pNPP), into the yellow para-nitrophenol (pNP), leading to an increase in absorbance of the light with wavelength of 405 nm (A405). The addition of chaperone Spy, Hsp70 A8, TkHsp20 and ClpB at a molar ratio of 5:1 relative to PHPT1 enhanced the efficiency of pNPP cleavage by PHPT1. As shown in Fig. 4B, the increase of A405 occurred more rapidly in all experimental groups of PHPT1 in the presence of chaperone Spy, Hsp70 A8, TkHsp20 and ClpB respectively, demonstrating an enhanced catalytic activity of PHPT1 by different chaperones (Fig. S11A). *Pfu* pol is an enzyme involved in DNA replication, which is widely used in biotechnology and diagnostic industries. To investigate the effect of chaperone on the *Pfu* pol, steady-state kinetic data was collected and analyzed using an EvaGreen-based fluorometric polymerase activity assay. The fit of the enzyme kinetic data with the Michaelis-Menten equation showed a significant improvement in *k*_cat_/*K*_m_, driven by the decrease in *K*_m_ (Fig. S11B), and represented a 1.75-fold improvement in catalytic activity with TkHsp20 (Fig. 4C). Xylanase is an enzyme widely used in the biofuel, paper, and food industries for breakdown of xylan. To evaluate the effect of chaperones on the activity of xylanase, 0.1 μM xylanase was utilized for the activity assay with xylan substrate varied from 0 mg/ml to 5 mg/ml in the presence or absence of molecular chaperones TkHsp20, Hsp70 A8, and ClpB respectively. The xylanase activity was measured by determining the release of reducing sugars from beechwood xylan according to the dinitrosalicylic acid (DNS) method (*33, 34*). The addition of each of chaperone TkHsp20, Hsp70 A8, and ClpB at a 10:1 molar ratio relative to xylanase enhanced the efficiency of xylan hydrolysis by xylanase arranging from 1.19-fold (Hsp70 A8) to 1.73-fold (ClpB) (Fig. 4D). Cas12a/Cas13a are gene editing enzymes with wide applications in gene editing, disease diagnostics, and synthetic biology applications. The trans-cleavage activity of Cas12a was investigated with a dsDNA substrate and ssDNA reporter labeled with a fluorophore (FAM) and a quencher (BHQ) in the presence and absence of chaperones ClpB, Hsp70 A8, Spy and TkHsp20, respectively. As shown in Fig. 4F, we observed a significant increase in Cas12a cleavage activity on the reporter upon the addition each of ClpB, Hsp70 A8, Spy and TkHsp20. Especially, the addition of TkHsp20 can increase the detection limit of Cas12a by up to 1.91-fold (Fig. S11C). Similarly, we investigated the trans-cleavage activity of Cas13a with an RNA substrate and ssRNA reporter labeled with a fluorophore (FAM) and a quencher (BHQ) in the presence and absence of 0.25 ng/μL TkHsp20. Fig. 4E showed a 1.43-fold improvement in catalytic activity with TkHsp20, and we observed a 1.23-fold increase in the detection limit of Cas13a (Fig. S11D). Collectively, these results indicated that molecular chaperones played a general role in modulating the activity of a wide range of enzymes.

**Figure 4.**
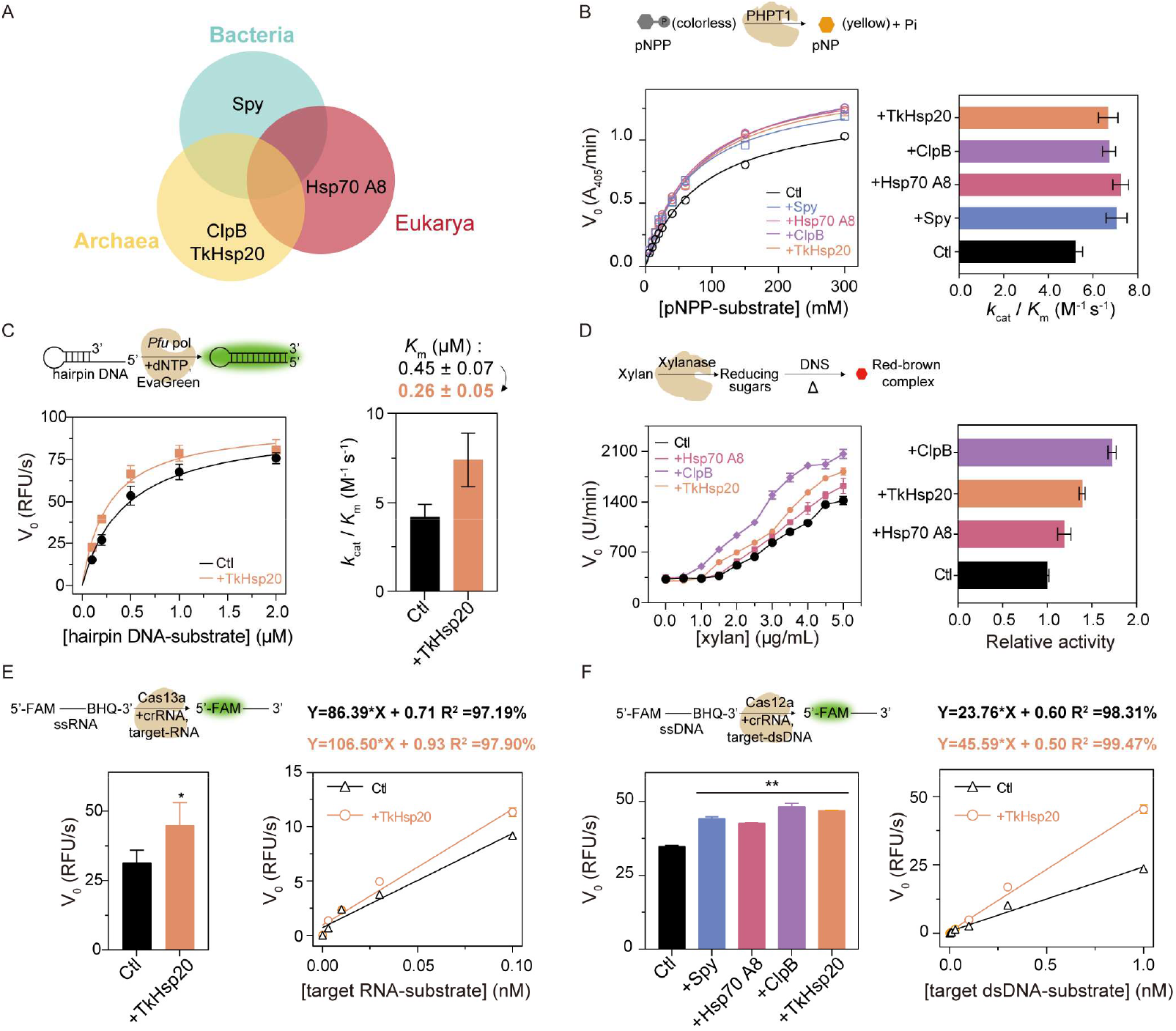
Molecular chaperones enhance the catalytic activity of various enzymes. (A) Species type of molecular chaperones used in the enzyme activity assays. (B) Substrate concentration-dependent enzymatic activity assays of PHPT1 in the absence and presence of Spy, Hsp70 A8, ClpB, and TkHsp20 respectively. Kinetic parameters *k*_cat_/*K*_m_ were calculated and compared for PHPT1 in the absence and presence of different chaperones. (C) Substrate concentration-dependent enzymatic activity assays of *Pfu* pol in the absence and presence of TkHsp20. Kinetic parameters *k*_cat_/*K*_m_ were calculated and compared for *Pfu* pol in the absence and presence of the chaperone TkHsp20. (D) Substrate concentration–dependent activity assays of xylanase performed in the absence or presence of the molecular chaperones Hsp70 A8, ClpB, and TkHsp20. The catalytic efficiency of xylanase under each condition was quantified relative to that of xylanase alone and normalized accordingly. (E) Enzymatic activity assay of Cas13a in the absence and presence of TkHsp20. (F) Enzymatic activity assays of Cas12a in the absence and presence of molecular chaperones Spy, Hsp70 A8, ClpB, and TkHsp20 respectively. All data represent the mean ± SD of three technical replicates.

## Discussion

Overall, our data showed chaperones from different categories influence the activity of the various enzymes. Chaperone Spy and Hsp70 A8, modulate the conformational exchanges of HEWL, stabilizing HEWL in its excited conformation, thereby enhancing its enzymatic activity. The mechanism by which chaperone Spy modifies the conformations of HEWL is of interest for other chaperone-enzyme interaction systems, as they may use similar mechanisms to modulate conformational exchanges of enzymes and regulate their activities. Both ATP dependent and independent chaperones are shown to recover misfolded client protein into their native states (*35-38*). This process of refolding of client proteins involves overcoming a free energy barrier to convert stable misfolded proteins into a different transiently unfolded or partially unfolded states with higher free energy, which, upon release, may spontaneously fold into its native state with lower free energy. From an energy landscape perspective, the denaturation effect of chaperones on misfolded client protein is similar to Spy’s modulation of HEWL conformational equilibrium. As in both processes, the folding energy barriers of client protein was reduced to ease the transition between different states (Fig. 5). The native states of most enzymes are marginally stable, with folding free energies typically ranging from 5 to 15 kcal mol^−1^, about the same amount of energy as 1∼3 hydrogen bond (*39-41*). As a result, enzymes can adopt alternative conformations within biologically relevant timescales. The energy landscape of an enzyme defines the number of thermodynamically and kinetically accessible conformations, their relative probabilities, and the rates at which they interconvert (Fig. 5). Thus, the energy landscape plays a crucial role in determining the functions of enzymes. Any perturbation of an enzyme or change of its environmental condition can alter the energy landscape and the conformational equilibrium by either lowering or raising energy barriers. The intrinsically dynamic properties of client enzymes making them more likely to adopt a conformation that chaperones can recognize. Chaperones then modulate the conformational exchanges of their client enzymes to either assist the client enzyme to adopt a conformation suitable for substrates binding or accelerating the convert rate of substrates. This is also in align with previous studies of conformational dynamics influence catalysis came from experiments by Tsou’s group (*42, 43*). They made the striking observation that small amounts of denaturant accelerated enzymatic function. This involvement of the entire energy landscape in catalysis was further suggested by the observation that applying and releasing a stretching force on an enzyme in a single molecular basis could increase the likelihood of enzymatic activity at a specific moment. Force-induced transition of the enzyme showed significant impact on the reactivity of chemical reactions and the alteration of the protein folding energy landscape (*44-46*). Therefore, the modulation of the protein folding free energy landscape and subsequently conformational equilibrium may represent a general mechanism employed by chaperones to modulate the activities of enzymes.

**Figure 5.**
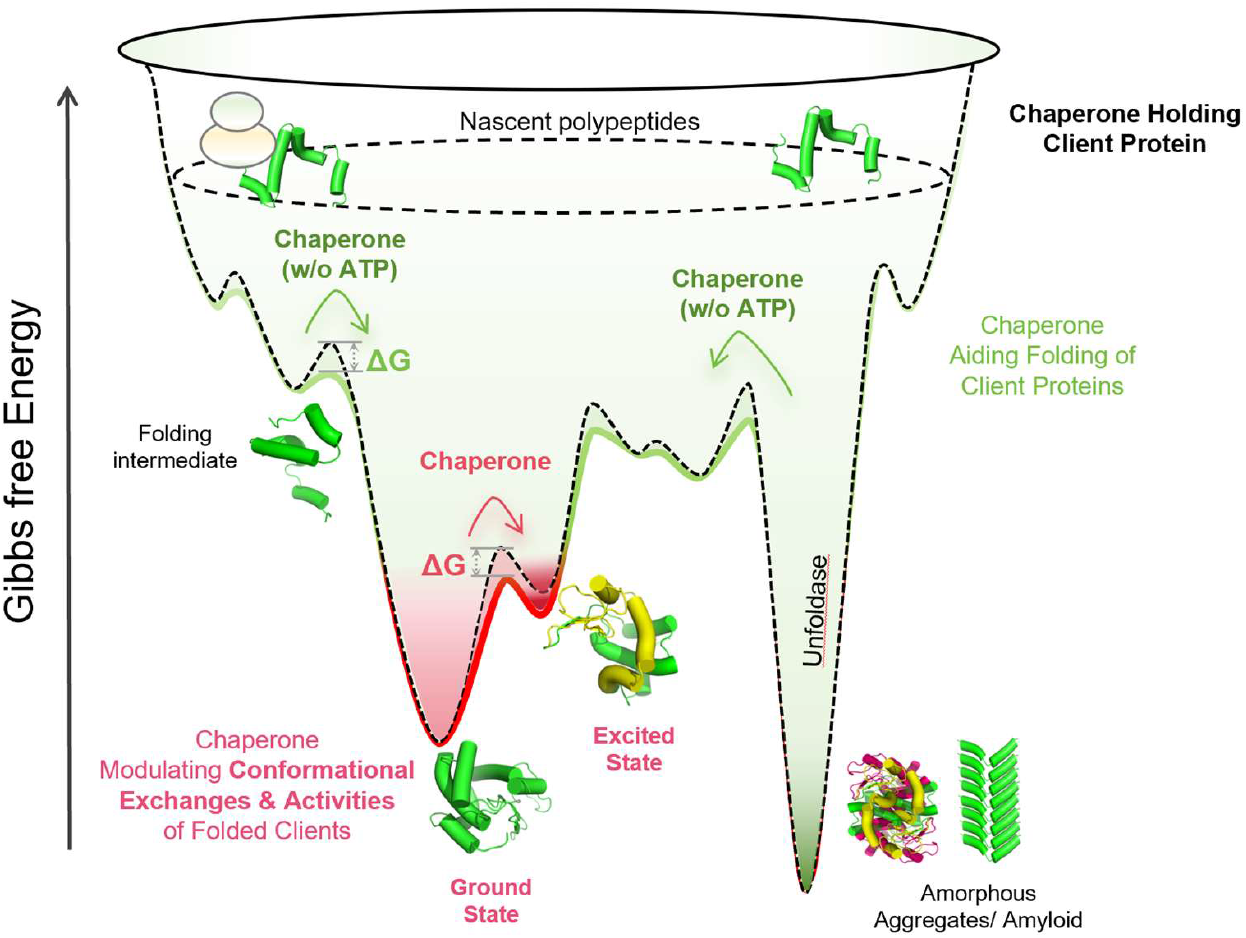
The energy landscape of chaperone’s function in maintain protein hemostasis and modulating the conformational exchanges of folded enzymes. Chaperones modify the ruggedness of the folding free energy landscape of client proteins by flattening the folding free energy landscape barrier. Folding intermediate of client proteins with high free energy are stabilized via interaction with chaperones. Amorphous and amyloid proteins with lower energy are destabilized and shifted towards the native state through hydrolyzing of ATP or entropic pulling. For folded client proteins, chaperones lower the energy barrier between ground and excited states.

To fully understand the role of chaperones in folded proteins, it is also crucial to know how they recognize client enzymes with diverse sizes and structures. Our previous research and others work demonstrated that four distinct chaperones—Spy, SurA, Skp and GroEL—bind to the same frustrated surface on folded client proteins, regardless of their sequence or structure (*13, 47*). Frustration arises when the constraints on amino acid residue connectivity conflict with their tendency to minimize free energy through interactions. This occurs in specific protein regions where the sequence cannot simultaneously achieve the optimal energy configuration for each residue. Consequently, these locally frustrated sites become energetically unstable, leading to increased flexibility and dynamic behavior. This local frustration site often corresponds to functional sites, underlying the dynamic interconverting conformational states of enzymes. The recognition and perturbation of the dynamics of local frustration sites will result in direct modification of the activity of enzymes. As we have observed in the interaction of HEWL with Spy and Hsp70 A8, both chaperones modulate the dynamics of the local frustrated site of HEWL in the region from residue 100 to 112, leading to the shift of the conformational exchanges of HEWL and subsequently its bacteriolytic activity.

Along with these mechanistic roles of chaperones in modulating enzyme conformational dynamics and activities, our findings underscore the potential usage of chaperones as auxiliary tools in enzyme-based applications. In diagnostics, the ability of chaperones to enhance catalytic efficiency and lower detection limits, as demonstrated for *Pfu* pol, CRISPR-Cas12a and Cas13a systems (Fig. 4C, E-F), suggests that chaperones could be exploited to improve the sensitivity of molecular diagnostic, including those used in pathogen detection and genetic screening. In biomanufacturing, chaperones could serve as additives to boost the activity and stability of industrial enzymes, such as the xylanase, thereby increasing yield and reducing production costs in processes of synthesis. The dynamics regulation mechanism of chaperones on modulating enzyme activities leads to precise adjustment in enzyme activities, offering exceptional tunability in enzyme-based biological processes.

Moreover, our results also suggest a new role of chaperones could play in physiological conditions to maintain the functional protein homeostasis. Chaperone proteins are among the most abundant in human cells, constituting as much as 10% of the total protein mass in these cells (*48, 49*). They are not only located in the cytoplasm but also in the nucleus, where no nascent poly peptide translation occurs (*8, 50-52*). The function of chaperones could be different in the cytoplasm and the nucleus. An intriguing possibility is that certain enzymes deliberately leverage their association with the chaperones as a regulatory mechanism. Unlike mechanisms operating at the levels of signal transduction to changing the level of transcription, translation, or degradation, the intrinsic dynamics of enzymes enable another way of regulation of the activity of enzymes by associating upregulated level of chaperones, allowing for greater adaptability to the environmental stress. Considering our observation that the enhancement of population of excited states and activities of enzymes depending on the concentration of chaperones (Fig. 1D, Fig. S2). Cells could regulate the functions of enzymes through the expression level of chaperones. Especially, for cancer cells or cells infected by pathogens, the chaperone expression level was upregulated up to 5-10 times (*53-55*), the excited states of enzymes could exhibit distinct activities under conformational modulation by interacted with various upregulated chaperones in cell.

In summary, our study uncovers the influence of chaperones on conformational exchanges and activities of folded enzymes. This reveals a new dimension to chaperone function, highlighting their broad potential not only in maintaining of proteostasis but also in enhancing enzyme-based applications across diagnostics, biomanufacturing and regulating protein functional homeostasis physiologically.

## Funding

This work is supported by

Strategic Priority Research Program of the Chinese Academy of Sciences XDB0540000

Natural Science Foundation of China grants 22327901, 22174151 and 21991080

Hubei Provincial Natural Science Foundation of China 2023AFA041

Postdoctoral Fellowship Program of China Postdoctoral Science Foundation GZC20232755

Hubei Postdoctoral Innovation Post Foundation of China R23R000304

## Author contributions

Conceptualization: L.H., B.Y., G.W., Y.C., X.Z., S.H., X.Z., M.L.; Methodology: L.H., B.Y., G.W., Y.C., S.Q., Z.G.; Investigation: G.W., Y.C., B.Y., R.D., S.Q.; Visualization: G.W., R.D., B.Y., Y.C.; Funding acquisition: L.H., M.L., X.Z., G.W.; Project administration: L.H.; Supervision: L.H.; Writing – original draft: B.Y.; Writing – review & editing: L.H., B.Y., G.W., Y.C.

## Data and materials availability

All data are available in the main text or the supplementary materials.

