## Supplementary Information for "A Regulatory Role of Molecular Chaperones in Excited States and Functions of Folded Enzymes"

Submitted Manuscript: Confidential
Template revised June, 2025

### Supplementary Materials for

#### An Unexpected Role of Molecular Chaperones in Regulating the Excited States and Activities of Folded Enzymes

Guan Wang[1†], Bin Yu[1†], Yihao Chen[1†], Ruishen Ding[1,2], Shixing Qi[1,2], Xiaoling Zhao[3,4], Zhou Gong[1,2], Xu Zhang[1,2], Sebastian Hiller[5], Maili Liu[1,2,6], Lichun He[1,2*]

**Affiliations:**

[1]State Key Laboratory of Magnetic Resonance Spectroscopy and Imaging, Innovation Academy for Precision Measurement Science and Technology, Chinese Academy of Sciences, Wuhan, Hubei, 430071, China.

[2]University of Chinese Academy of Sciences, Beijing, 100049, China.

[3]Department of Reproductive Medicine, General Hospital of Central Theater Command of the People's Liberation Army, Wuhan, Hubei 430061, China.

[4]Qinhe Life Science Ltd. Wuhan 430000, China.

[5]Biozentrum, University of Basel, 4056, Basel, Switzerland.

[6]Optics Valley Laboratory, Hubei 430074, China.

[†]These authors contributed equally to this work.

**The PDF file includes:**

Materials and Methods
Figs. S1 to S11
Tables S1
References

### Supplementary Materials

#### Materials and Methods

##### Preparation of protein

The pPIC9K-XynCDBFV vector was synthesized by Huamei Biological Engineering Inc. (Wuhan), and the *HEWL* gene was codon-optimized for *P pastoris* expression. The pPIC9K-XynCDBFV-HEWL plasmid was linearized with BglII (TAKARA, Japan) and transformed into *P. pastoris* GS115 by electroporation. Positive transformants were selected on histidine-deficient MD plates and cultured in 1L YPD medium at 30°C and 220 rpm. When the OD600 reached ~30, cells were harvested by centrifugation (5,000 xg, 10 min, 25°C) and resuspended in 1L FM22 medium supplemented with 2% glycerol. After 2 days of cultivation, cells were transferred to FM22 medium containing 2g $^{15}NH_4Cl$ and 1% methanol, with additional 1% methanol added every 24 h for the protein induction. Following 3 days of induction, the supernatant was collected and concentrated by tangential flow filtration. The sample was sequentially purified using SP column and Superdex 75 column to obtain $^{15}$N-labeled HEWL according to a published protocol (*1*).

The gene of *PHPT1* and *HEWL E35Q* mutant were synthesized by Sangon Biotech Inc. and cloned into the pET21a vector, containing an N-terminal, TEV-cleavable His6-SUMO-tag. The plasmids encoding the molecular chaperones Spy, Spy cysteine mutants, Hsp70 A8, TkHsp20, and ClpB are stored in our laboratory. For the expression of molecular chaperone and PHPT1, the plasmid was transformed into the T7 *E.coli* strain and cultured in the TB medium containing 100 µg/mL ampicillin or 50 µg/mL kanamycin at 37°C and 220 rpm. When the OD600 reached ~2, 0.5 mM Isopropyl β-D-1 thiogalactopyranoside (IPTG) was added to induce protein expression at 25°C for 6 h. Spy and its mutants were purified as described previously (*2*). For purification of Hsp70 A8, TkHsp20, ClpB, PHPT1, and HEWL E35Q mutant, cells were resuspended in PBS buffer (20 mM Sodium Phosphate, 300 mM NaCl, pH 7.4), and lysed by high-pressure homogenization. Following centrifugation (20,000 ×g, 1 h, 25°C) to remove debris, the proteins were purified by Ni-NTA gravity columns. Elution fractions containing the target proteins were collected and dialyzed against PBS buffer in the presence of SUMO protease to facilitate overnight cleavage. The digest was then applied to a reverse Ni-NTA gravity flow column to remove cleaved tags and contaminants. Subsequently, the protein was concentrated and further purified by size-exclusion chromatography using a Superdex 75 column to obtain the purified protein.

##### Assay of PHPT1 activity

The enzymatic activity of PHPT1 was assayed in a 100 µL reaction mixture containing 50 mM Tris-HCl (pH 7.2) and 10 mM DTT at 37°C. To evaluate the effect of chaperones on the activity of PHPT1, chaperones were supplemented into the reaction mixture at a molar ratio of 5:1 relative to PHPT1. Kinetic analysis was performed over 1 h using 7.1 µM PHPT1 with the substrate para-nitrophenyl phosphate (pNPP) at concentrations of 6, 10, 15, 20, 25, 40, 60, 150, and 300 mM. The concentration of the product, para-nitrophenol (pNP) was determined by measuring absorbance at 405 nm. The initial reaction rate for each template concentration was determined by calculating the first derivative of the substrate absorbance curve at 405 nm. Kinetic parameters ($k_{cat}$ and $K_m$) were derived by fitting the Michaelis-Menten equation to the initial rate data using GraphPad Prism 8.0.

##### Assay of xylanase activity

Xylanase (0.1 µM) was utilized for activity assay with xylan substrate varied from 0 mg/ml to 5 mg/ml in the presence or absence of molecular chaperones Hsp70 A8, Spy, and ClpB. The xylanase activity was assayed by determining the release of reducing sugars from beechwood xylan according to the dinitrosalicylic acid (DNS) method (*3*). To evaluate the effect of chaperones on the activity of xylanase, chaperones were supplemented into the reaction mixture at a molar ratio of 10:1 relative to xylanase. Each sample was incubated in a 37°C water bath for 30 min at pH5.5. After 30 min, 0.5 mL of DNS was added to stop the reaction, and the mixture was then boiled for 5 min. The A540 nm absorbance value was measured with a multi-functional microplate reader. The average reaction rate was determined at 30 minutes across varying substrate concentrations. One unit (U) of enzyme activity is defined as the amount of enzyme that released 1 μM of reducing sugars per minute under the given conditions.

**Assay of HEWL activity**

The bacteriolytic activity of the recombinant HEWL against *M. lysodeikticus* was determined according to the standard spectrophotometry method (*4*). 40 μL HEWL (5 ug/mL) was mixed with 200 μL cell suspension with OD450 at 1.0 ± 0.1 in 20 mM phosphate buffer (20 mM sodium phosphate, 50 mM NaCl, pH 6.0). A decrease in the OD value at 450 nm was recorded every 1 min for 2 h, and the average reaction rate at 10 minutes was used to calculate the antibacterial activity of HEWL. The amount of enzyme decreasing the OD at 450 nm by 0.001 - per minute was defined as one U of enzyme activity.

**Assay of Pfu polymerase activity**

Steady-state kinetic data were collected and analyzed using an EvaGreen-based fluorometric polymerase activity assay (*5*). A hairpin DNA template (5'-CCAGCATTATGAAAGTGACACG TGCACCATTGGTGCACGTG-3') was prepared at 100 μM in water by heating at 98°C for 5 min followed by annealing on ice for 30 min. For kinetic measurements, varying concentrations of the hairpin DNA template (0.1-2 μM) were mixed with dNTPs and 8 nM *Pfu* DNA polymerase in reaction buffer (25 mM Tris-HCl, 10 mM KCl, 0.05% Triton X-100, pH 8.8) in the presence and absence of 0.25 ng/μL TkHsp20. Reactions were initiated at 72°C by adding $MgCl_2$ to a final concentration of 2.5 mM, with dNTPs maintained at 200 μM throughout. Fluorescence was monitored over time, and background signals from control reactions were subtracted to generate background-corrected fluorescence curves. The initial reaction rate for each template concentration was determined by calculating the first derivative of the fluorescence curve. Kinetic parameters ($k_{cat}$ and $K_m$) were derived by fitting the Michaelis-Menten equation to the initial rate data using GraphPad Prism 8.0. Reported values represent the mean of biological duplicates, each comprising three technical replicates.

**Cas12a-based cleavage assay**

Cas12a-based cleavage assay was referenced a new method (*6*). Double-stranded DNA (dsDNA) activator was formed by annealing NTS (5'-AATAGGTGATTTTGGTCTAGCTACAGAGAA ATCTCGATG-3') and TS (5'-CATCGAGATTTCTCTGTAGCTAGACCAAAATCACCTA TT-3') primers in a 1:1 molar ratio by 95°C for 5 min followed by cooling to 37°C for 30 min. 2 μL of 100 nM *Lachnospiraceae bacterium* Cas12a (LbaCas12a was obtained from New England Biolabs, Inc), 2 μL of 200 nM crRNA (5'-UAAUUUCUACUAAGUGUAGAUGUCUAG CUACAGAGAAA-3'), 2 μL of 10× NEBuffer r2.1, and 10 μL of RNase-free $H_2O$ were added to make the total volume 16 μL and incubated for 15 min at 37°C in the presence and absence of different chaperones (0.25 ng/μL TkHsp20 or 0.5 ng/μL ClpB or 0.5 ng/μL Hsp70 A8 or 0.25 ng/μL Spy). Then, 2 μL of 10 nM dsDNA activator and 2 μL of 2.5 μM ssDNA reporter (5'6-FAM-TTTTTTTTTTTTTTT-3'BHQ1) were added to a final volume of 20 μL. The fluorescence intensity was recorded immediately every 30 s at 37°C. For Limit of Detection (LOD) of the

Cas12a measurements, 2 μL of 100 nM Cas12a, 2 μL of 200 nM crRNA, 2 μL of 10× NEBuffer r2.1 and 10 μL of RNase-free $H_2O$ were added and incubated for 15 min at 37°C in the presence and absence of 0.25 ng/μL TkHsp20. Then, 2 μL of dsDNA activator and 2 μL of 2.5 μM ssDNA reporter were added. The fluorescence intensity was measured as mentioned above. The concentration of dsDNA activator was 3, 10, 30, 100, 300, and 1000 pM for ssDNA reporter in NEBuffer r2.1. The LOD was calculated using the 3σ/K method.

**Cas13a-based cleavage assay**

Typical Cas13a-based cleavage assay was described previously (*7*). Briefly, The *Leptotrichia wadei* Cas13a (LwaCas13a was obtained from Magigen company) mediated cleavage assays were carried out using the reaction buffer (20 mM HEPES, 50 mM NaCl, 6 mM $MgCl_2$, pH 7.5). The experimental procedure included the following steps. Initially, 1 μL of 1 μM LwaCas13a, 1 μL of 1 μM crRNA (5'-GAUUUAGACUACCCCAAAAACGAAGGGGACUAAAACACACU ACCUGCACUAUAAGCACUUUAGUGC-3'), 1 μL of 40 U/μL RNase inhibitor and 17 μL of the reaction buffer were mixed and preincubated at 37°C for 10 min in the presence and absence of 0.25 ng/μL TkHsp20. After the formation of the LwaCas13a-crRNA complex, 10 μL of 10 nM target RNA (5'-GUAGCACUAAAGUGCUUAUAGUGCAGGUAGUGUUUA-3') and 20 μL of 1 μM ssRNA reporter (5'6-FAM-mAAUGGCmA-3'BHQ1) were added to create a 50 μL reaction solution. The fluorescence intensity was recorded immediately every 30 s at 37°C. For LOD of the Cas13a measurements, 1 μL of 100 nM Cas13a, 1 μL of 200 nM crRNA, 1 μL of 40 U/μL RNase inhibitor, and 17 μL of reaction buffer were added and incubated for 10 min at 37°C in the presence and absence of 0.25 ng/μL TkHsp20. Then, 10 μL of target RNA and 20 μL of 1 μM ssRNA reporter were added. The fluorescence intensity was measured as mentioned above. The concentration of RNA activator was 0, 3, 10, 30, and 100 pM for ssRNA reporter in reaction buffer. The LOD was calculated using the 3σ/K method.

**Solvent PRE**

For solvent PRE measurements of both HEWL and Spy-HEWL complex, two corresponding samples, one without and one with 2 mM Gd-DOTA (SIGMA-ALDRISH), were prepared in 20 mM phosphate buffer and measured with identical experimental parameters on Bruker Avance-600 spectrometers at 25°C. Relaxation delay $d_1$ was set to 8 s for sufficient relaxation before each scan. Peak intensities were obtained with the software CcpNmr (*8*). The solvent PRE shielding index, $\sigma_i$, was introduced as the following to quantify the shielding effect:

$$\sigma_i = \left(\frac{I_i^{Spy}}{I_0^{Spy}} - \frac{I_i^{buffer}}{I_0^{buffer}}\right) \Big/ \left(\frac{I_i^{buffer}}{I_0^{buffer}}\right) \qquad (1)$$

where $I_0^{Spy}$ is the intensity of HEWL residues in the presence of Spy without TEMPOL, $I_0^{buffer}$ is the intensity of HEWL residues in the buffer without TEMPOL. $I_i^{Spy}$ is the intensity of HEWL residues in the presence of Spy with TEMPOL and $I_i^{buffer}$ is the intensity of HEWL residues in the buffer with TEMPOL.

**Site-specific PRE experiments**

Spin labeling of the cysteine mutants of Spy with MTSL(Toronto Research Chemicals) was done according to a published protocol (*9*). PD-10 columns (GE Healthcare) were used to remove the DTT before linking with MTSL and to remove the excess of the MTSL after labeling. To reduce the spin label, a final concentration of 5 mM ascorbate was added to the NMR tube from a 500 mM pH-adjusted stock solution. NMR samples contained 0.15 mM $^{15}$N-labeled HEWL and approximately 0.4 mM unlabeled Spy. Two-dimensional (2D) [$^{15}$N,$^{1}$H]-HSQC NMR spectra were recorded before and after addition of sodium ascorbate at 25°C using an 600 MHz spectrometer equipped with a cryogenic probe. Intermolecular paramagnetic relaxation effect on $^{15}N$-labeled HEWL upon interaction with the spin label MTSL-attached T35C, M53C, T99C and T124C mutants of Spy, calculated as $I_{ox}/I_{red}$, the ratio of the heights of peaks before and after the reduction of the spin label.

**H/D exchange measurements**

A 500 μL sample of 1 mM $^{15}N$-labeled HEWL or a complex sample containing 1 mM $^{15}N$-labeled HEWL and 1.6 mM unlabeled Spy were buffer-exchanged into an identical buffer with 90% $D_2O$ using NP-5 desalting columns. Consecutive 2D [$^{15}N$,$^{1}H$]-SFHMQC spectra for each sample were acquired on a Bruker Avance 600 MHz spectrometer at 25°C. The first [$^{15}N$,$^{1}H$]-SFHMQC spectra was acquired for each sample starting precisely 10 min after $D_2O$ buffer exchange. Relaxation delay $d_1$ was set to 0.1 s. Peak intensities were obtained with the software CcpNmr (*8*).

In the consecutive 2D HSQC measurements of the H/D exchange rate, the exchange rate $k_i$ of the client protein residue i was fitted with equations (3) and (4):

$$I_{ti} = I'_{0i}e^{-k_i t} + I'_{0i}a \qquad (2)$$

$$I'_{0i} = \frac{I_{0i}}{0.9} \qquad (3)$$

where $t$ was the exchange time; $I_{0i}$ was the peak intensity of residue i at time zero with 90% $D_2O$. $I'_{0i}$ was the back calculated peak intensity of residue i at time zero with 100% $D_2O$; $I_{ti}$ was the peak intensity of residue $i$ at time t; $k_i$ was the exchange rate of residue $i$; $a$ was the percentage of $H_2O$.

Calculations of the hydrogen exchange protection factor and Gibbs free energy of exchange were referred to a published work[7] , the hydrogen exchange protection factor (PF) of residue $i$ was calculated with the formula:

$$PF_i = \frac{k_{i_{intr}}}{k_{i_{ex}}} \qquad (4)$$

where $k_{i_{intr}}$ represents the intrinsic chemical rate of residue $i$ in a fully open state at a given pH, temperature. The protection factor $PPPP_{ii}$ was related to the opening free energy via the formula (5) :

$$\Delta G = RTIn^{PF-1} \qquad (5)$$

**Spectral density functions**

To determine the longitudinal and transverse relaxation rates of HEWL, $T_1$, $T_2$, and {$^{1}H$}-$^{15}N$ NOE data were collected. $T_1$ data were acquired with relaxation delays of 30 ms, 50 ms, 100 ms, 200 ms, 300 ms, 400 ms, 500 ms, 700 ms, 900 ms, and 1100 ms. $T_2$ data were acquired with relaxation delays of 16.34 ms, 32.68 ms, 49.02 ms, 65.36 ms, 81.7 ms, 98.04 ms, 114.38 ms, 130.72 ms, 147.06 ms, 163.4 ms, 179.74 ms and 196.08 ms. All NMR spectra were processed with the software NMRPipe (*10*) and analyzed with the software CcpNmr (*8*). The reduced spectral density mapping was employed to characterize the backbone dynamics of apo HEWL and HEWL in the Spy-HEWL complex according to a published protocol (*11*). The average value for each plot is indicated by a horizontal reference line.

**Backbone $^{15}N$ CPMG RD experiments**

For the micro- to mili-second time-scale chemical exchange measurement, $^{15}N$ Carr-Purcell-Meiboom-Gill relaxation dispersion (CPMG RD) experiments were recorded in an interleaved manner at two different magnetic fields, 600 MHz and 800 MHz (*12*). A constant delay of 40 ms was used with a series of CPMG field strengths, defined as $\nu_{cpmg} = 1/2t_{cp}$ (50, 100, 150, 200, 250, 300, 350, 400, 450, 500, 550, 600, 650, 700, 750, 800, 900, 1000 Hz) where $t_{cp}$ is the delay between two consecutive 180 pulses, and two repeat experiments were carried on at $\nu_{cpmg}$ = 100 and 500 Hz for the error estimation. Effective relaxation rate, $R_{2,eff}$ was calculated according to the following equation:

$$R_{2,eff} = -\frac{1}{T_{CPMG}} In\left(\frac{I_{\nu_{CPMG}}}{I_0}\right) \qquad (6)$$

where $T_{CPMG}$ is the constant time delay, $I_0$ and $I_{\nu_{CPMG}}$ are peak intensities in the absence and presence of a CPMG field, respectively. The relaxation dispersion data was analyzed by fitting to a two-site exchange model using the software ChemEx (https://github.com/gbouvignies/ChemEx). Uncertainties in $R_{2,eff}$ were calculated as:

$$\Delta R_{2,eff} = \frac{1}{T_{CPMG}} \frac{\Delta I}{I_{\nu_{CPMG}}} \qquad (7)$$

where $\Delta I$ is the average standard deviation of peak intensities estimated from the repeat measurements (*13*).

**Backbone $^{15}N$ CEST experiments**

$^{15}N$ CEST experiments were carried out at 25°C on a Bruker Avance 600-MHz spectrometer. Pseudo-3D spectra were acquired with the $^{15}N$ carrier frequencies positioned from 100 ppm to 132 ppm at a spacing of 0.5 ppm during the irradiation time of $T_{EX}$ = 500 ms. Irradiation field strengths $B_1$ of 30 Hz was used for HEWL. All the spectra were processed with the NMRPipe program. The CEST profile for each individual residue was generated by calculating the intensity ratios $I_i/I_{i0}$ versus the varied $^{15}N$ carrier frequencies, where $I_{i0}$ is the intensity of residue i measured in the reference spectrum, and $I_i$ is the intensity of residue i measured with the application of the $B_1$ field. For residues showing minor dip, the data were fitted to a two-state exchange model with the python program ChemEx (https://github.com/gbouvignies/ChemEx).

**Isothermal titration calorimetry measurements**

Experiments were carried out on the PEAQ-ITC calorimeter (MicroCal). The designed cell volume and syringe volume for PEAQ-ITC were 212.7 µL and 40 µL respectively. Protein samples were dialyzed to the same buffer (20mM sodium phosphate, 50mM NaCl, pH6) at 4°C overnight prior to experiments. Protein concentrations were determined after dialysis. Experiments were performed at 25°C with an initial injection of 0.4 µL followed by a series of injections of 2 µL.

**Figs. S1 to S11**

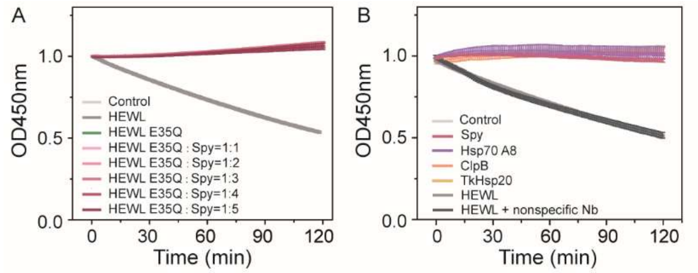


**Figure S1. Enzyme activity analysis.** (A) Time-dependent enzymatic activity of E35Q mutant in the presence of molecular chaperones Spy at molar ratios of 1:1, 2:1, 3:1, 4:1, and 5:1 relative to E35Q. (B) Time-dependent enzymatic activity of molecular chaperones Spy, Hsp70 A8, ClpB and TkHsp20. Molecular chaperones do not possess catalytic activity. In the presence of only molecular chaperones Hsp70 A8, ClpB, Spy, and TkHsp20, no hydrolysis of the substrate *M. lysodeikticus occurs*.

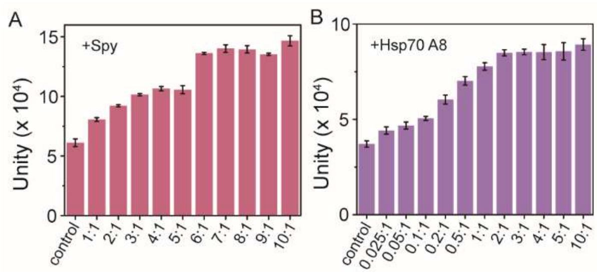


**Fig S2. The enhancement of HEWL activity by molecular chaperones Spy and Hsp70 A8 is concentration-dependent.** (A) Statistical analysis of HEWL catalytic activity in the presence of different concentrations of molecular chaperone Spy. (B) Statistical analysis of HEWL catalytic activity in the presence of different concentrations of molecular chaperone Hsp70 A8.

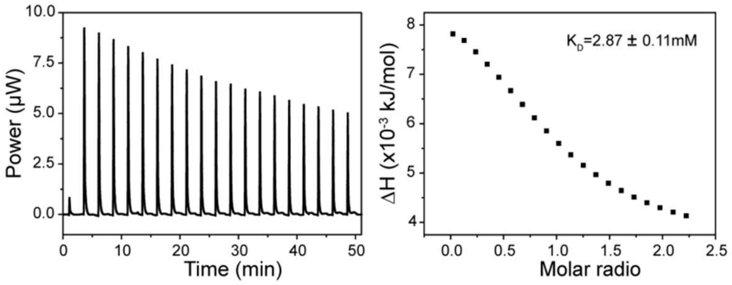


**Fig S3. ITC measurement of interactions between Spy and HEWL.** Titration of 81.34 mg/mL (5650 µM) HEWL injected in steps of 2 µL to a cell volume of 212.7 µL containing 16.02 mg/mL (504 µM Spy). The x-axis indicates the titrant to titrate molar ratio. The peak volume of each injection is integrated with Affinimeter (*14*).

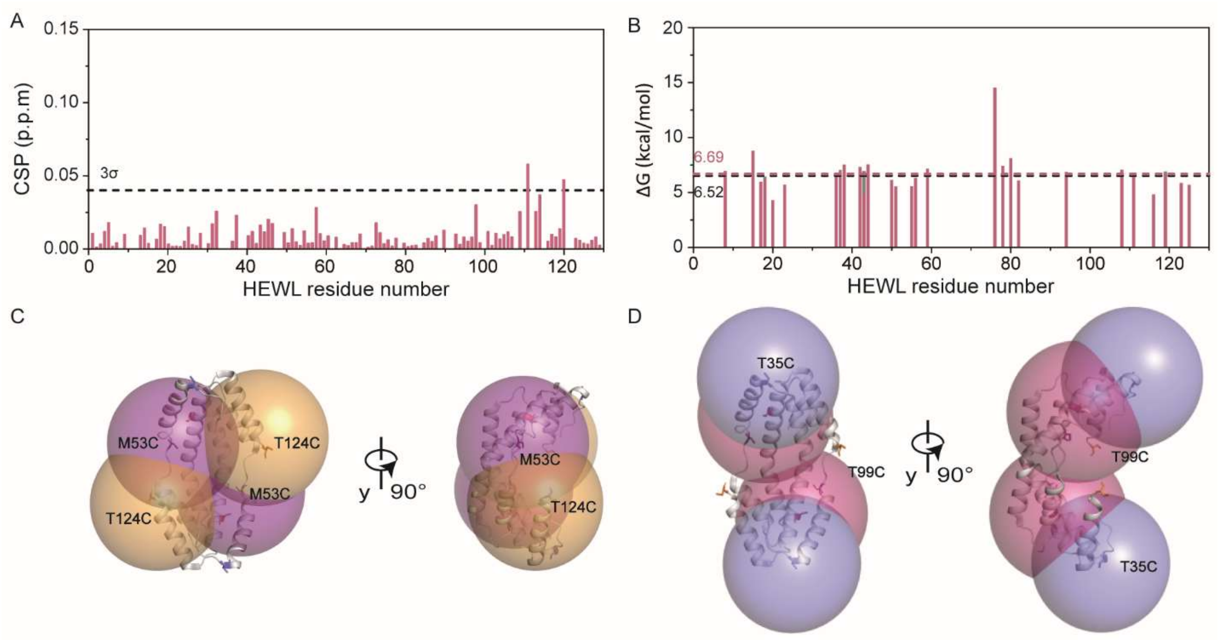


**Fig S4. Molecular chaperone Spy interacts with HEWL in a non-specific and transient manner.** (A) CSPs of amide moieties of 1 mM $^{15}$N-labeled HEWL plotted against the corresponding amino acid residue number after adding equimolar concentrations of molecular chaperones Spy. (B) The free energy of exchange of backbone amides of ~0.7 mM HEWL in the absence (grey) and presence (red) of ~1 mM Spy are plotted against the amino acid residue number. (C-D) The PRE effect range, visualized as spheres with a 20 Å radius, are displayed on the crystal structure of Spy (PDB 3O39). For each spin label, spheres are centered around the $C_\beta$ atom of each cysteine mutant of Spy.

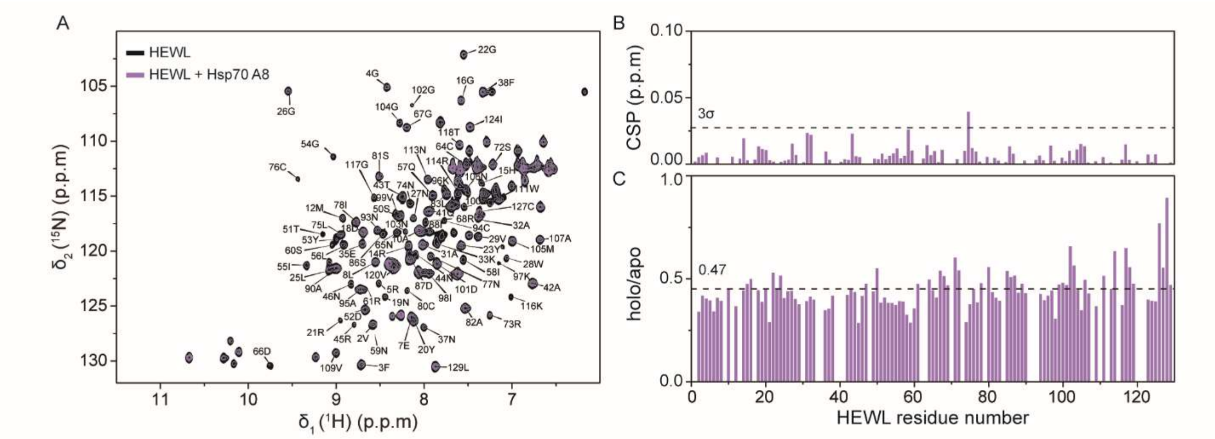


**Fig S5. Molecular chaperone Hsp70 A8 interacts with HEWL in a non-specific and transient manner.** (A) Overlay of [$^{15}$N,$^{1}$H]-HSQC spectra of HEWL in the absence (black) and presence (purple) of Hsp70 A8. (B) CSPs of amide moieties of $^{15}$N-labeled HEWL after adding Hsp70 A8. (C) Ratios of the residual resolved cross peak volume of backbone amides of 1 mM $^{15}$N-labeled HEWL in the presence and absence of 0.05mM Hsp70 A8 plotted against the corresponding amino acid residue number.

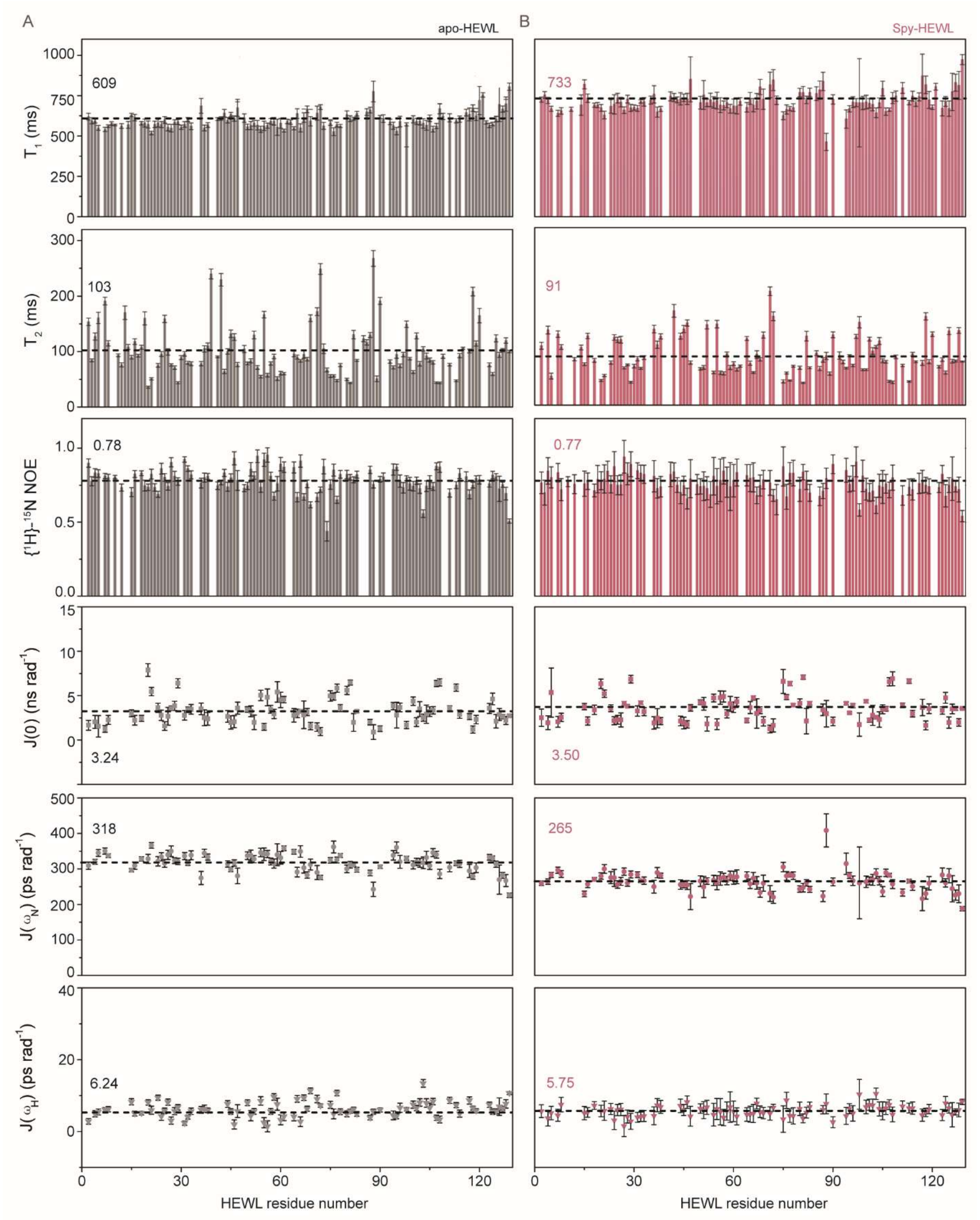


**Fig S6. Amplitudes of backbone motions of HEWL.** Backbone motions of HEWL were measured by the relaxation parameters ($T_1$, $T_2$, Hetero-NOE) and spectral density function, in its apo- state (A) and holo- state with Spy (B) at 25℃**.** Dashed lines indicate the sequence-averaged value.

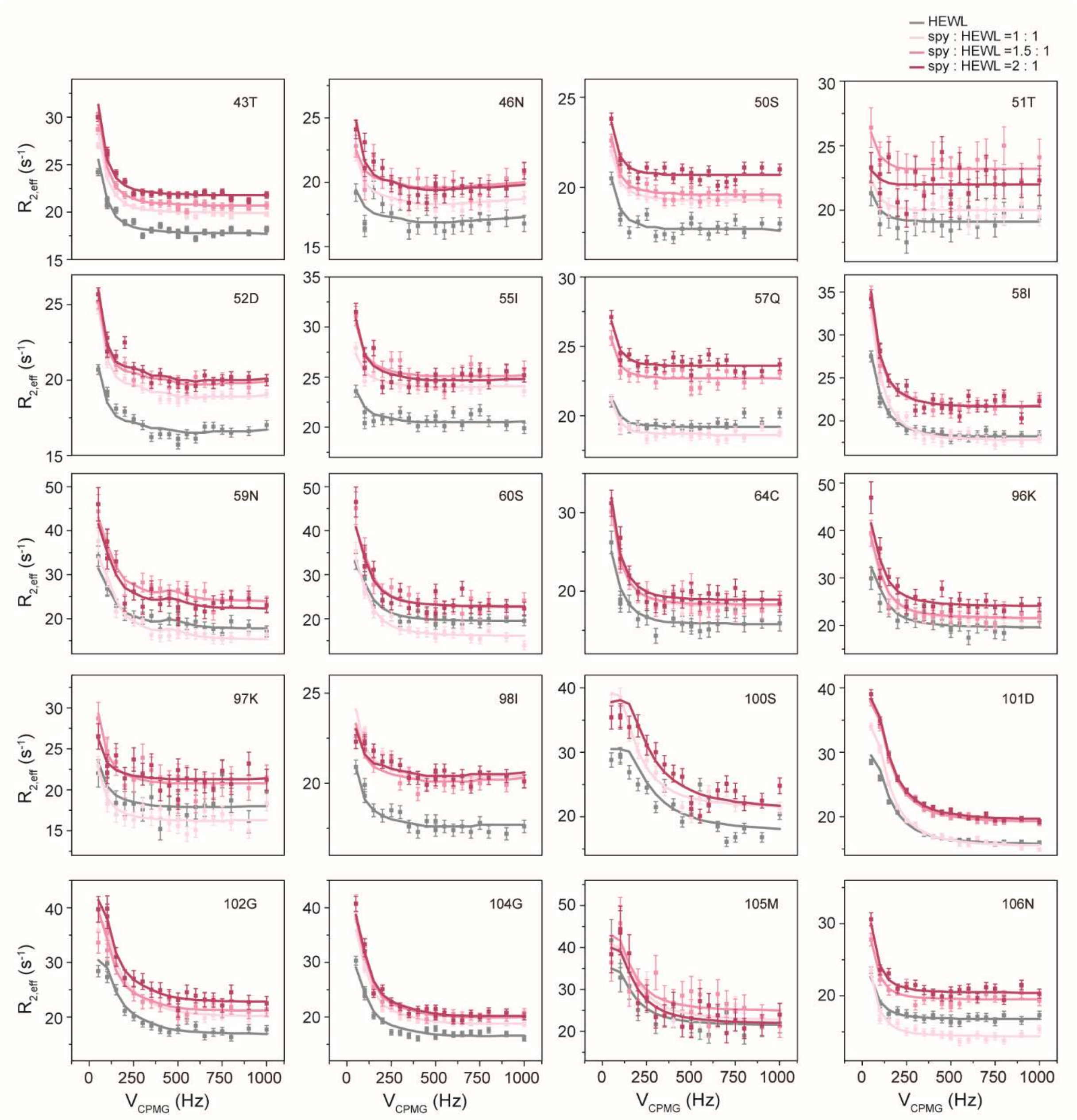


**Fig S7. Relaxation dispersion profiles of HEWL.** Global fitting of $^{15}$N CPMG RD profiles for HEWL in the absence and presence of molecular chaperones Spy at molar ratios of 1:1, 1.5:1, and 2:1 relative to HEWL. Data were collected at $^{1}$H frequencies of 600MHz at 25℃.

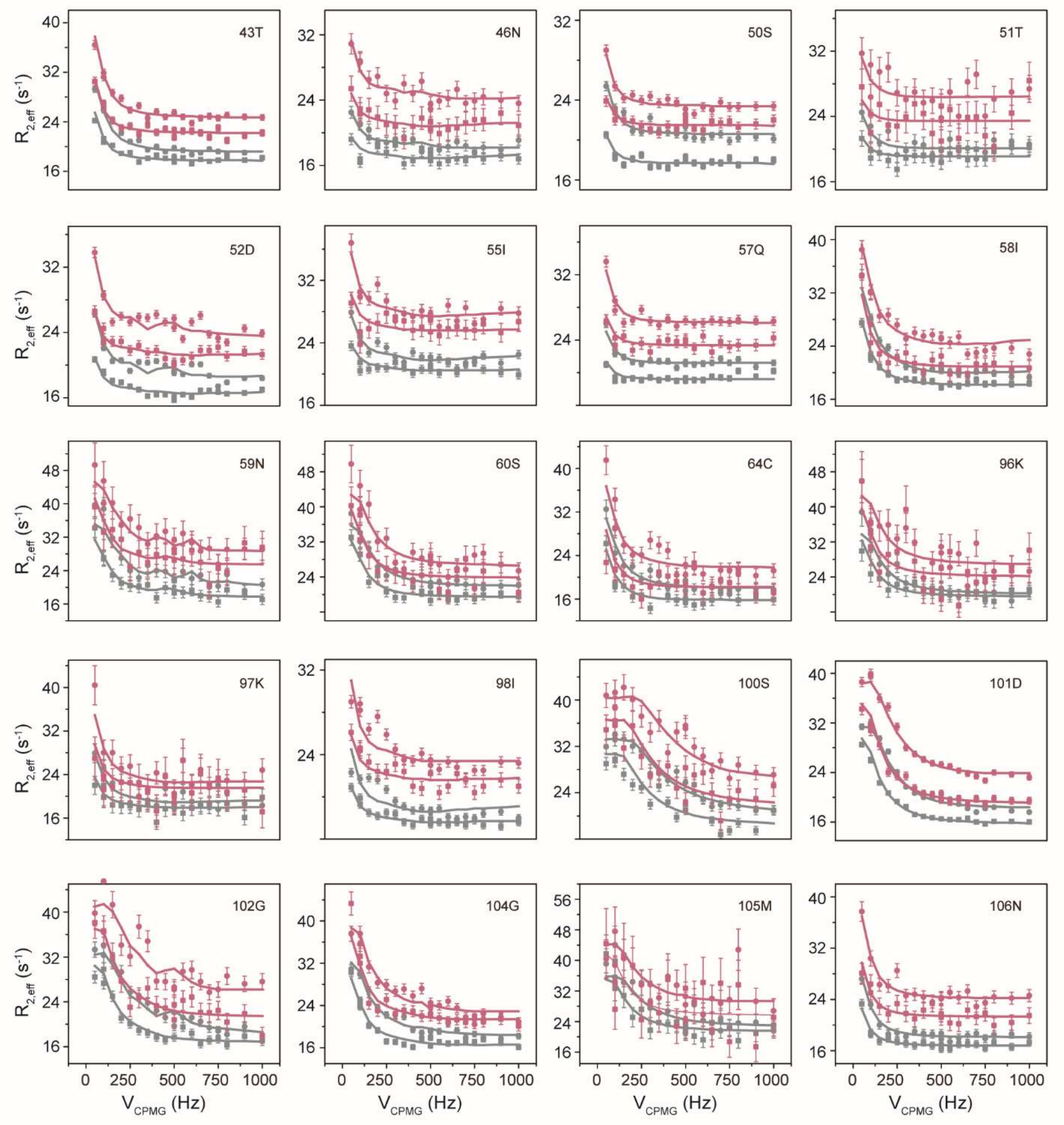


**Fig S8. Relaxation dispersion profiles of HEWL.** Global fitting of $^{15}N$ CPMG RD profiles for HEWL in the absence (black) and presence (purple) of Spy at molar ratio of 2:1. Data were collected at $^{1}H$ frequencies of 600 (square) and 800MHz (circle) respectively.

5

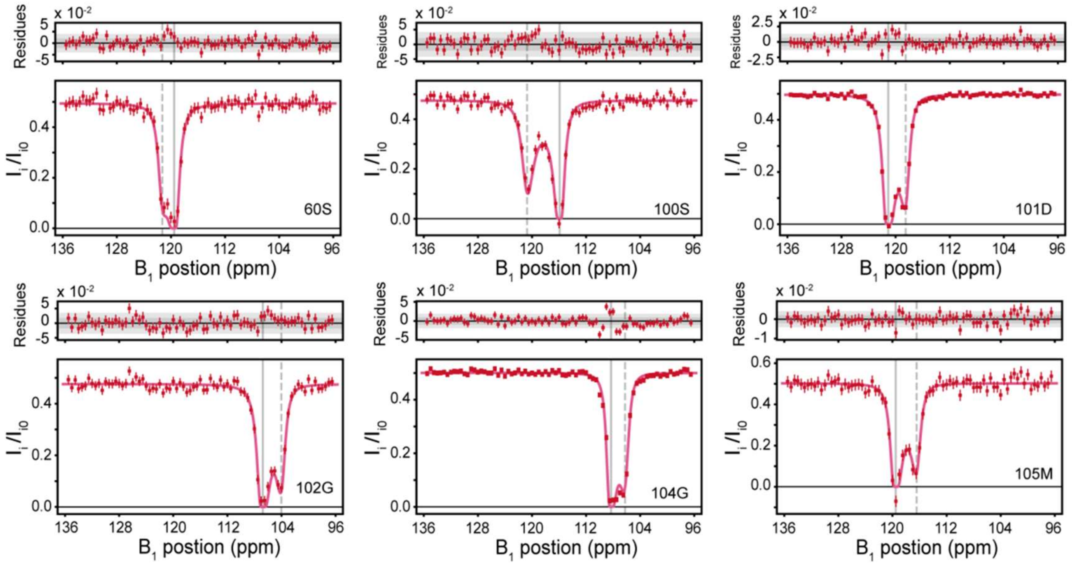


**Fig S9. $^{15}$N CEST profiles recorded with a $B_1$ field of 30 Hz for residues 60S, 100S, 101D, 102G, 104G and 105M of HEWL in the buffer.** Each profile graphs the ratios ($I_i/I_{i0}$) of peak intensities in the presence ($I_i$) and absence ($I_{i0}$) of a weak $B_1$ field as a function of the position where the $B_1$ field is applied. The large and small dips represent the chemical shifts of the corresponding residues in different conformations.

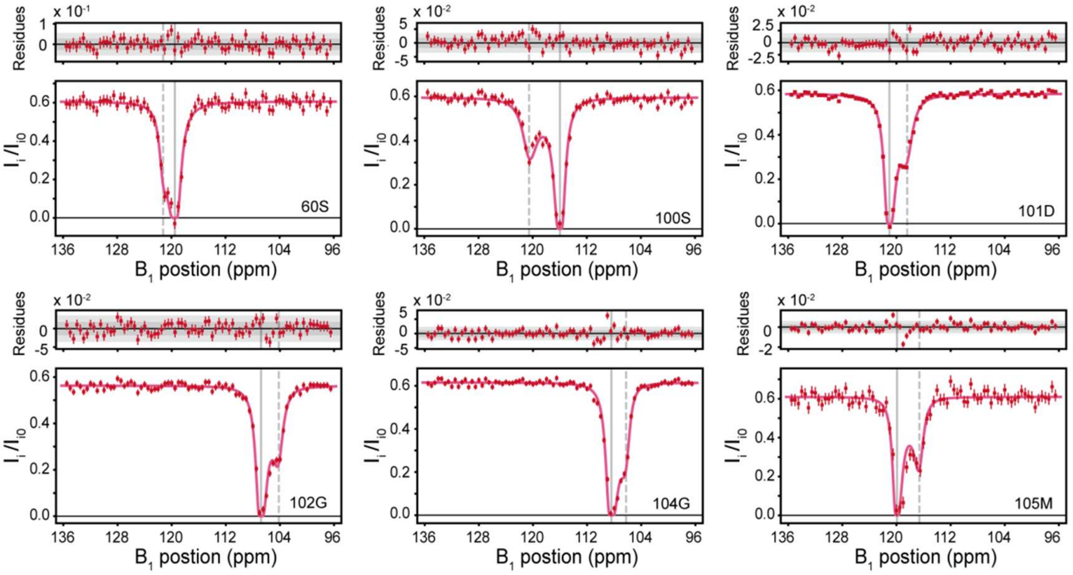


**Fig S10. $^{15}$N CEST profiles recorded with a $B_1$ field of 30 Hz for residues 60S, 100S, 101D, 102G, 104G and 105M of HEWL in the presence of molecular chaperones Spy at molar ratios of 2:1 relative to HEWL at $^1$H frequencies of 600MHz.** Each profile graphs the ratios ($I_i/I_{i0}$) of peak intensities in the presence ($I_i$) and absence ($I_{i0}$) of a weak $B_1$ field as a function of the position where the $B_1$ field is applied. The large and small dips represent the chemical shifts of the corresponding residues in different conformations.

A

| | $k_{cat}$ ($min^{-1}$) | $K_m$ (mM) | $k_{cat}$ / $K_m$ ($M^{-1}s^{-1}$) |
|---|---|---|---|
| PHPT1 apo | 2.58 ± 0.06 | 82.50 ± 4.77 | (5.21 ± 0.32) × $10^{-1}$ |
| PHPT1 + Spy | 2.86 ± 0.07 | 67.49 ± 4.13 | (7.06 ± 0.47) × $10^{-1}$ |
| PHPT1 + Hsp70 A8 | 3.05 ± 0.06 | 70.17 ± 3.12 | (7.24 ± 0.35) × $10^{-1}$ |
| PHPT1 + ClpB | 3.05 ± 0.05 | 75.64 ± 3.04 | (6.72 ± 0.29) × $10^{-1}$ |
| PHPT1 + TkHsp20 | 3.16 ± 0.08 | 78.78 ± 4.67 | (6.68 ± 0.43) × $10^{-1}$ |

B

| | $k_{cat}$ ($S^{-1}$) | $K_m$ (μM) | $k_{cat}$ / $K_m$ ($M^{-1}s^{-1}$) |
|---|---|---|---|
| *Pfu* pol apo | 190.70 ± 11.07 | 0.45 ± 0.07 | (4.2 ± 0.7) ×$10^8$ |
| *Pfu pol* + TkHsp20 | 192.68 ± 11.20 | 0.26 ± 0.05 | (7.4 ± 1.5) ×$10^8$ |

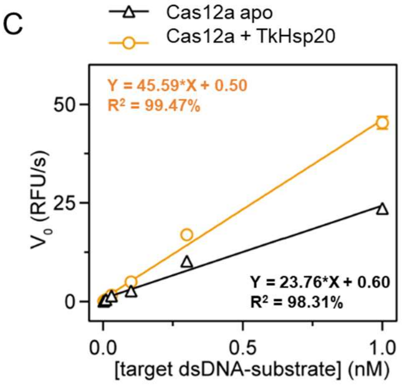


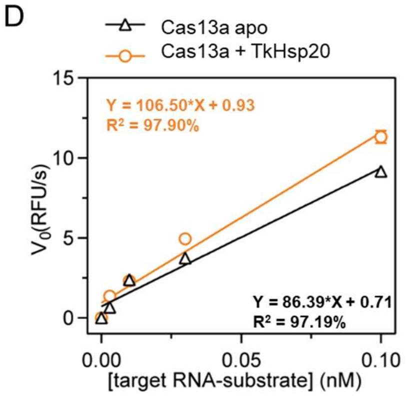


**Fig S11. Molecular chaperones enhance the catalytic activity of different enzymes.** (A) Kinetic parameters for *Pfu* pol in the absence and presence of TkHsp20. (B) Kinetic parameters for PHPT1 in the absence and presence of different chaperones. (C) Detection limits of Cas12a in the presence and absence of TkHsp20. (D) Detection limits of Cas13a in the presence and absence of TkHsp20.

**Table S1. Exchange Parameters for HEWL.**

| Client HEWL | $k_1$ ($s^{-1}$) | $k_{-1}$ ($s^{-1}$) | $k_{ex}$ ($s^{-1}$) | $P_E$ (%) |
|---|---|---|---|---|
| apo[1] | 17.9 ± 0.5 | 256 ± 21 | 274 ± 22 | 6.53 ± 0.45 |
| Spy :HEWL=1:1 | 22.2 ± 1.0 | 277 ± 22 | 299 ± 23 | 7.42 ± 0.37 |
| Spy:HEWL=1.5:1 | 21.6 ± 1.2 | 257 ± 26 | 279 ± 28 | 7.75 ± 0.54 |
| Spy :HEWL=2:1 | 20.4 ± 0.9 | 211 ± 25 | 231 ± 26 | 8.81 ± 0.86 |
| apo[2] | 17.6 ± 1.3 | 190 ± 40 | 208 ± 41 | 8.45 ± 1.41 |
| A8:HEWL=1:20 | 29.1 ± 2.9 | 171 ± 26 | 200 ± 30 | 14.52 ± 1.62 |
| E35Q | -- | -- | -- | -- |
| Spy:E35Q=2:1 | -- | -- | -- | -- |

-- Data not available.
[1] HEWL is used as a control for Spy titration in 20 mM Sodium Phosphate Buffer, 50 mM NaCl, pH 6.0.
[2] HEWL is used as a control for Hsp70 A8 titration in 20 mM Sodium Phosphate Buffer, 50 mM NaCl, pH 7.4.

**References (*1–14*)**
